# Prostaglandin EP3 Receptor signaling is required to prevent insulin hypersecretion and metabolic dysfunction in a non-obese mouse model of insulin resistance

**DOI:** 10.1101/671289

**Authors:** Jaclyn A. Wisinski, Austin Reuter, Darby C. Peter, Michael D. Schaid, Rachel J. Fenske, Michelle E. Kimple

**Author notes:** To whom correspondence should be addressed: Michelle E. Kimple, PhD, 4148 UW Medical Foundation Centennial Building, 1685 Highland Avenue, Madison, WI 53705, OR Jaclyn A. Wisinski, PhD, 3031 Cowley Hall, 1725 State Street, LaCrosse, WI 54601.

## Abstract

When homozygous for the *Leptin*^*Ob*^ mutation (Ob), Black-and-Tan Brachyury (BTBR) mice become morbidly obese and severely insulin resistant, and by 10 weeks of age, frankly diabetic. Previous work has shown Prostaglandin EP3 Receptor (EP3) expression and activity is up-regulated in islets from BTBR-Ob mice as compared to lean controls, actively contributing to their beta-cell dysfunction. In this work, we aimed to test the impact of beta-cell-specific EP3 loss on the BTBR-Ob phenotype by crossing *Ptger3* floxed mice with the Rat insulin promoter (RIP)-Cre^*Herr*^ driver strain. Instead, germline recombination of the floxed allele in the founder mouse – an event whose prevalence we identified as directly associated with underlying insulin resistance of the background strain – generated a full-body knockout. Full-body EP3 loss provided no diabetes protection to BTBR-Ob mice, but, unexpectedly, significantly worsened BTBR-lean insulin resistance and glucose tolerance. This *in vivo* phenotype was not associated with changes in beta-cell fractional area or markers of beta-cell replication *ex vivo*. Instead, EP3-null BTBR-lean islets had essentially uncontrolled insulin hypersecretion. The selective up-regulation of constitutively-active EP3 splice variants in islets from young, lean BTBR mice as compared to C57BL/6J, where no phenotype of EP3 loss has been observed, provides a potential explanation for the hypersecretion phenotype. In support of this, high islet EP3 expression in Balb/c females vs. Balb/c males was fully consistent with their sexually-dimorphic metabolic phenotype after loss of EP3-coupled Gα_z_ protein. Taken together, our findings provide a new dimension to the understanding of EP3 as a critical brake on insulin secretion.

**New and Noteworthy:** Islet Prostaglandin EP3 receptor (EP3) signaling is well-known as up-regulated in the pathophysiological conditions of type 2 diabetes, contributing to beta-cell dysfunction. Unexpected findings in mouse models of non-obese insulin sensitivity and resistance provide a new dimension to our understanding of EP3 as a key modulator of insulin secretion. A previously-unknown relationship between mouse insulin resistance and the penetrance of Rat insulin promoter-driven germline floxed allele recombination is critical to consider when creating beta-cell-specific knockouts.

**For Table of Contents Use Only:** 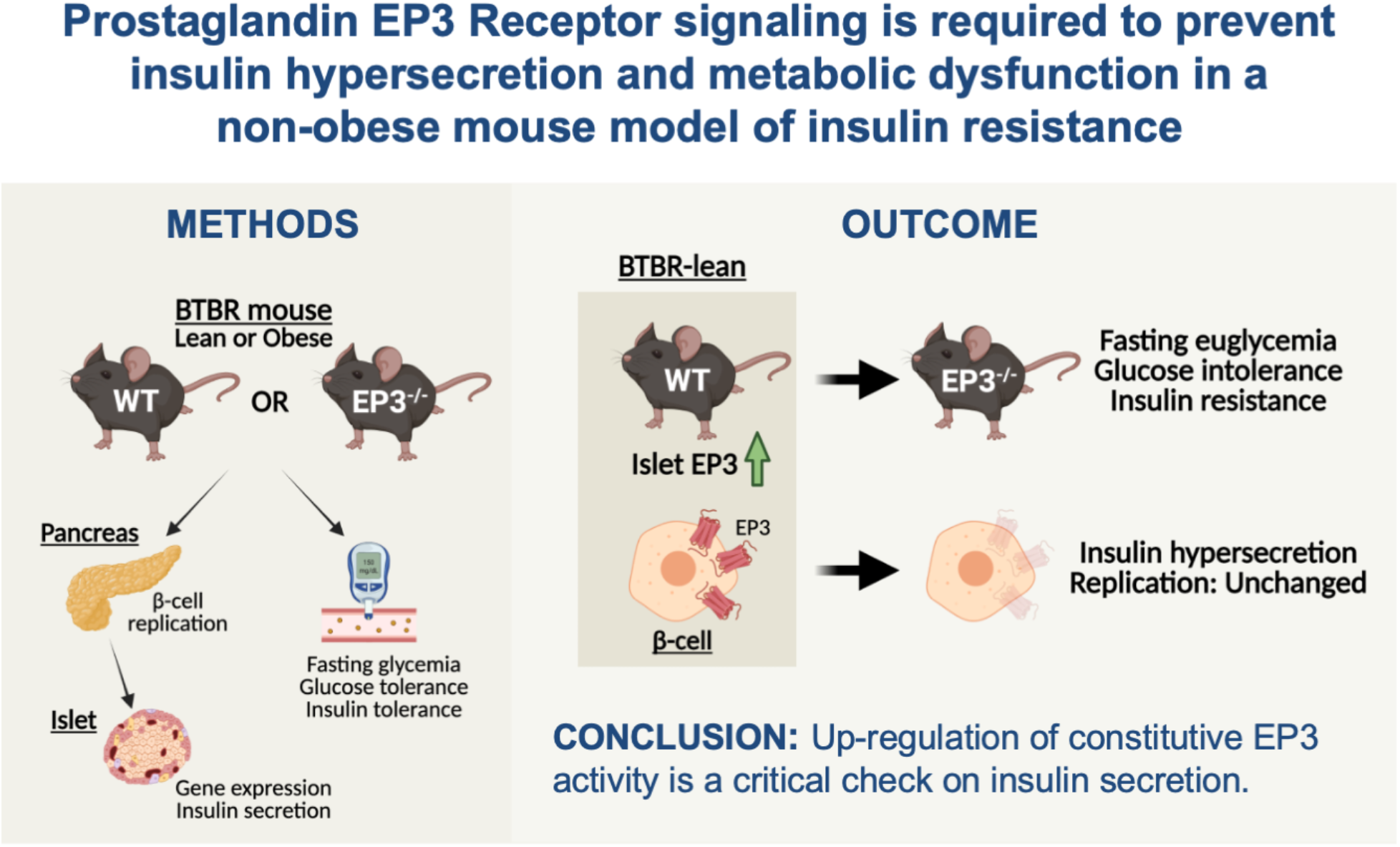

## Introduction

Type 2 diabetes (T2D) occurs after a failure of the insulin-secreting pancreatic beta-cells to compensate for peripheral insulin resistance and its often-associated glucolipotoxic and inflammatory conditions. Genetically-encoded defects in both beta-cell function or the ability of beta-cells to proliferate and survive can be factors in an individual’s susceptibility to the disease. As a strain characteristic, Black and Tan Brachyury (BTBR) mice are more insulin resistant than the more commonly used C57BL/6J sub-strain (1, 2), but their T2D pathogenesis when homozygous for the *Leptin*^*Ob*^ mutation (Ob) is beta-cell centric. The BTBR strain has an underlying deficit in insulin granule exocytosis (3, 4) and fails to up-regulate an islet cell cycle gene module necessary to mount a compensatory increase in beta-cell mass (5-7). Male and female BTBR-Ob mice rapidly and reproducibly develop severe obesity and insulin resistance, succumbing to T2D by 10 weeks of age (2, 8, 9). This makes the BTBR mouse an excellent model to mimic the natural course of T2D development in susceptible individuals.

Prostaglandin EP3 receptor (EP3) is a G_i/o_-coupled receptor for the arachidonic acid metabolite, prostaglandin E_2_ (PGE_2_): its most abundant natural ligand. When EP3 is active, adenylyl cyclase activity is inhibited and cAMP production reduced, blunting glucose-stimulated insulin secretion (GSIS) (8-15). Islets from BTBR-Ob mice and T2D human organ donors express more EP3 and produce more PGE_2_ than those of non-diabetic controls, and an EP3 antagonist can improve their GSIS response (8, 9, 13). EP3 antagonists have also been shown to increase mouse and human beta-cell proliferation (16, 17). Because of these findings, elevated beta-cell EP3 signaling has been proposed as a dysfunctional consequence of the T2D condition itself that could be therapeutically targeted *in vivo* (18-23). Arguing against this concept are studies showing (a) global EP3 loss magnifies the development of obesity, insulin resistance, and/or glucose intolerance of aging or after moderate-fat- or high-fat diet feeding (24-26) and (b) EP3 antagonism fails to improve the T2D phenotype of diet-induced obese mice (27). Therefore, the critical question of whether EP3 signaling is protective or detrimental towards T2D phenotype remained unanswered.

In this work, we used the insulin-resistant BTBR mouse strain, both lean and Ob, to explore the role of EP3 *in vivo* metabolic function. Our original intent was to create a beta-cell-specific EP3-null BTBR mouse line, but, due to germline Cre recombination, we ultimately created and validated our model as a full-body knockout. Nevertheless, with the strong link between BTBR-Ob beta-cell dysfunction and upregulated islet PGE_2_ production and EP3 expression, we anticipated EP3-null BTBR-Ob mice would be protected from T2D. Our strongest findings, though, were in the BTBR-lean background, and were associated with significantly elevated expression of constitutively-active EP3 splice variants in BTBR-lean islets as a strain characteristic. In concert with the previously unreported and sexually-dimorphic metabolic phenotype of male and female Balb/c mice, similarly associated with differential islet EP3 splice variant expression, these findings provide a new dimension to our understanding of EP3 as a critical brake on insulin secretion. Finally, our finding the penetrance of germline RIP-Cre recombination is associated with the underlying insulin resistance of the background strain serves as a cautionary note to other investigators using constitutive insulin-promoter Cre driver lines to generate beta-cell-specific knockout mice.

## Materials & Methods

### Chemicals and Reagents

Sodium chloride (S9888), potassium chloride (P3911), magnesium sulfate heptahydrate (M9397), potassium phosphate monobasic (P0662), sodium bicarbonate (S6014), HEPES (H3375), calcium chloride dehydrate (C3881), exendin-4 (E7144), and RIA-grade bovine serum albumin (A7888) were purchased from Sigma Aldrich (St. Louis, MO, USA). RPMI 1640 medium (11879–020: no glucose), penicillin/streptomycin (15070–063), and fetal bovine serum (12306C: qualified, heat-inactivated, USDA-approved regions) were from Life Technologies (Carlsbad, CA, USA). Dextrose (D14–500) was from Fisher Scientific (Waltham, MA).

### Mouse husbandry

Breeding colonies were housed in a limited-access, pathogen-free breeding core facility where all cages, enrichment, and water were sterilized before use on a 12-hour light/12-hour dark cycle with *ad libitum* access to water and irradiated breeder chow (Teklad 2919, Envigo). Upon weaning, mice were housed four or fewer per cage of mixed genotypes with ad libitum access to lower-fat chow in the Madison VA Animal Resource Facility (LabDiet 5001, nonirradiated). All procedures were performed according to approved protocols by the principles and guidelines established by the University of Wisconsin-Madison, and William S. Middleton Memorial Veterans Hospital Animal Care and Use Committees.

B6/129 mice in which the first coding exon of the *Ptger3* gene is flanked by LoxP sites (EP3-floxed) were obtained from The Jackson Laboratory (*Ptger3*^tm1Csml^; stock 008349, deposited by Michael Lazarus). These mice were crossed with C57BL/6J mice in which Cre expression is driven by the rat insulin promoter (RIP-Cre^*Herr*^: “InsPr-Cre” in (28)). As compared to other constitutive RIP-Cre driver lines, RIP-Cre^*Herr*^ has low hypothalamic Cre expression (29) and does not contain the human growth hormone minigene, which has been shown in previous studies to produce full-length, active hGH that artificially enhances mouse beta-cell proliferation through the prolactin receptor (30, 31). The mixed-background EP3-flox-RIP-Cre mouse was back-crossed into the BTBR *T+ Itpr3*^*tf*^/J (BTBR) background more than 10 generations. In order to transfer the line to the UW Madison Biomedical Research Model Services Breeding Core facility, embryos obtained from in vitro fertilization of wild-type BTBR oocytes with sperm from EP3-flox-RIP-Cre males were implanted into pseudopregnant BTBR females. Because of the poor results of in vitro fertilization in the BTBR strain (32), only four pups were born from two in vitro fertilization trials, with only one founder mouse containing the EP3-floxed allele. This founder mouse was used to re-derive the colony, after which experimental mice were transferred to our PI-accessible animal facility.

The generation, validation, and husbandry of Gα_z_-null mice in the Balb/c background has been previously described (33). For analysis of islet EP3 expression, 9-week-old male and female Balb/c mice were purchased from Jackson Laboratories and acclimated for 1 week prior to terminal islet isolation.

### Mouse metabolic phenotyping

For BTBR phenotyping experiments, the only statistically significant change in metabolic parameters-of-interest was a slightly lower body weight in 8-week female mice as compared to male (data not shown). Therefore, both male and female mice were used. Oral glucose tolerance tests (OGTTs) were performed at 6 weeks of age after a 4-6 h fast in clean cages. Glucose levels before and after gavage of 1 g/kg glucose, up to 2 h post-gavage, were quantified as previously described (9, 12, 22, 34). Insulin tolerance tests (ITTs) were performed similarly to OGTTs, except that 0.75 mg/kg Humulin-R (Eli Lilly and Co) was IP injected.

For Balb/c experiments, IP GTTs and ITTs were performed on 9-12-week-old male and female mice after a 12 h fast and 4-6 h fast, respectively, and data analyzed separately by sex, as described previously for female Balb/c mice (33). Plasma insulin levels at baseline and 5-7.5 min after glucose challenge were determined using the Crystal Chem Inc. high-sensitivity rat/mouse insulin ELISA as previously described (33, 35).

### Islet isolation and tissue collection

To isolate pancreatic islets, mice were anesthetized by 2,2,2 tribromoethanol until unresponsive, exsanguinated by cutting the descending aorta, and the bile duct ligated and cannulated, followed by injection of ice-cold collagenase solution to inflate the pancreas as described previously (36). Whole pancreases were harvested upon necropsy and fixed in formalin at 4 degrees C overnight, followed by paraffin embedding. Adipose tissue and kidney were harvested upon necropsy and flash-frozen in liquid nitrogen for future isolation of total RNA.

### Quantification of beta-cell fractional area and beta-cell replication

Insulin immunohistochemistry with hematoxylin counterstain (for measurement of beta-cell fractional area) was performed as previously described (11, 34, 35). Co-immunofluorescence experiments with anti-insulin and anti-Ki67 antibodies to quantify actively-replicating beta-cells were performed as previously described (11, 34, 35). Guinea pig anti-insulin (discontinued), background-reducing antibody diluent (S302281-2), EnVision™ diaminobenzidine (DAB) reagents, and serum-free protein block (X090930-2) were from Dako (Agilent). Rabbit anti-Ki67 mAb D3B5 (9129) was from Cell Signaling. Citrate based antigen retrieval solution (H-3300) and Vectashield Mounting Medium with DAPI (H-2000) were from Vector Laboratories. FITC-coupled donkey anti-rabbit antibody (711-095-152) and Cy3-coupled goat anti-guinea pig antibody (707-165-148) were from Jackson Immunoresearch.

### Ex vivo islet glucose-stimulated insulin secretion (GSIS) assays

Immediately after isolation, islets were plated into 96-well V-bottomed tissue-culture-treated plates and cultured overnight in RPMI medium with 8.4 mM glucose, as previously described (37). Single-islet microplate GSIS assays were performed and total islet insulin content and secreted insulin quantified via an in-house sandwich ELISA as previously described(37). Anti-insulin antibodies (Insulin + Proinsulin Antibody, 10R-I136a; Insulin + Proinsulin Antibody, biotinylated, 61E-I136bBT) were from Fitzgerald Industries (Acton, MA, USA). The 10 ng/ml insulin standard (8013-K) and assay buffer (AB-PHK) were from Millipore.

### Quantitative PCR assays

150-200 islets from each mouse islet preparation were washed with PBS and used to generate RNA samples via the RNeasy Mini Kit (Qiagen) according to the manufacturer’s protocol. Copy DNA (cDNA) was generated and relative qPCR performed via SYBR Green assay using primers validated to provide linear results upon increasing concentrations of cDNA template, as previously described(35). The RNeasy Mini Kit (74104) and RNase-free DNase set (79254) were from Qiagen. High-Capacity cDNA Reverse Transcription Kit (4368814) was from Applied Biosystems. FastStart Universal SYBR Green Master Mix (4913850001) was from Millipore-Sigma. A list of qPCR primer sequences can be found in **Table 1**.

**Table 1:**
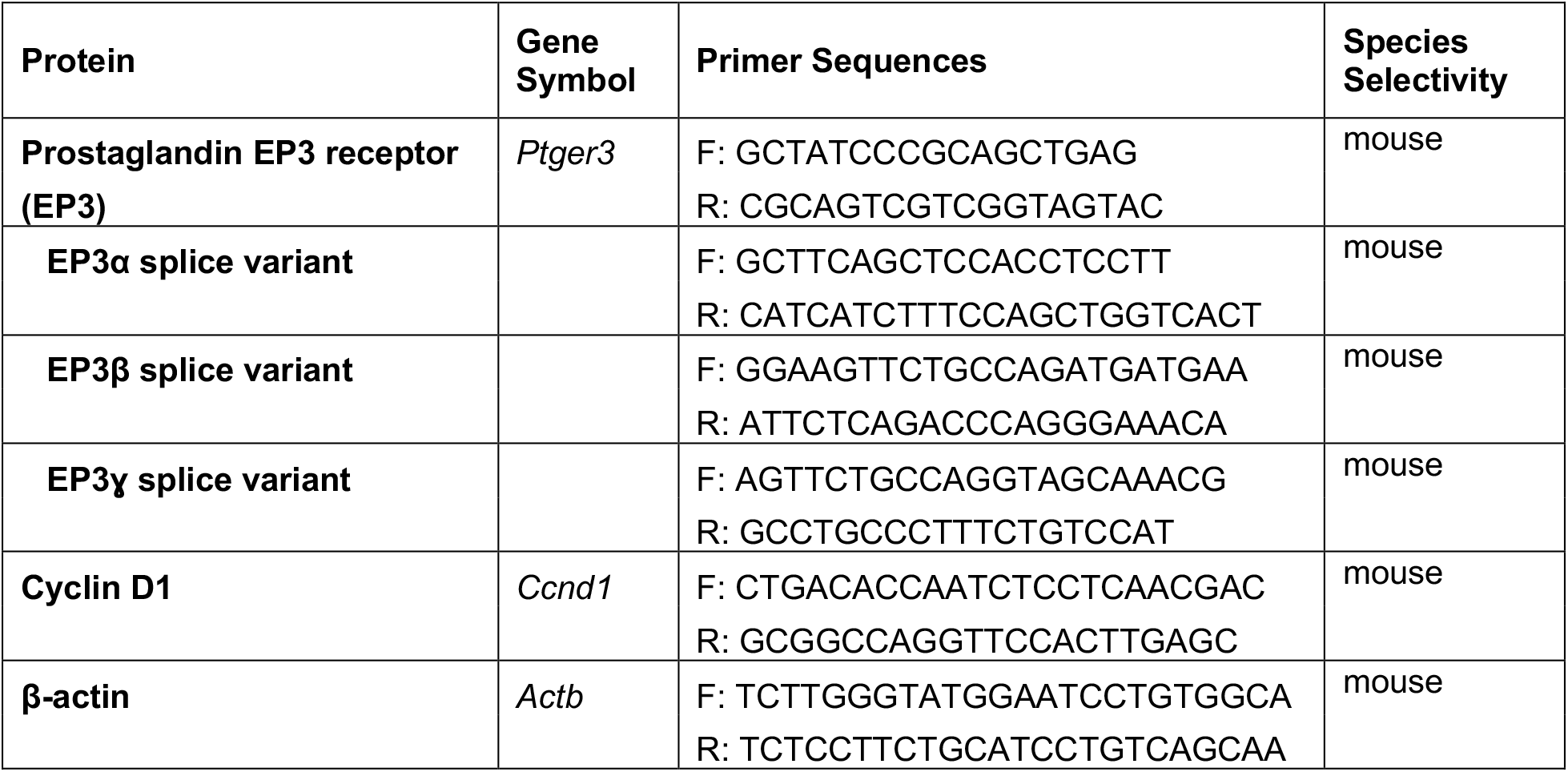
Quantitative PCR primer sequences.

### Statistical analysis

Data are expressed as mean ± standard error of the mean (SEM) unless otherwise noted. Data were analyzed as appropriate for the experimental design and as indicated in the figure legends. P-values < 0.05 were considered statistically significant. Statistical analyses were performed with GraphPad Prism version 9 (GraphPad Software, San Diego, CA).

## Results

### Germline recombination of the EP3-floxed allele in the RIP-Cre^*Herr*^ driver line created a full-body EP3-null mouse

As described in the Introduction, our original intent was to generate a beta-cell-specific EP3-null BTBR mouse. Upon NCBI BLAST search, we found the EP3-floxed genotyping primers recommended by The Jackson Laboratory amplify a 310 base pair product in both the intact and recombined (i.e., deleted) allele. Because of this, we designed two forward and one reverse primer that would give discrete bands for the WT, intact or deleted allele in genotyping reactions (**Table 2**). Supporting a potential full-body EP3 knockout, only the deleted allele was represented in archived ear punch samples from our colony, including the founder mouse, fully independent of Cre expression (**Figure 1A**). Adipose and kidney have the highest EP3 expression in the body, and no *Ptger3* mRNA was detected in adipose or kidney cDNA samples from EP3-flox-RIP-Cre mice, whether lean or obese (**Figure 1B,C**). Furthermore, the elevated EP3 expression in WT BTBR-Ob islets was completely ablated in EP3-flox-RIP-Cre islets, confirming a full-body knockout (EP3-null) (**Figure 1D**). While we were unable to determine the penetrance of EP3-Flox-RIP-Cre germline floxed allele recombination, extrapolating data from three different floxed protein-of-interest RIP-Cre^*Herr*^ mouse colonies revealed a strong association with the underlying insulin resistance of the background strain. Deleted alleles were detected over 5.4% of the time in samples from insulin-resistant BTBR mice, 3.9% percent of the time samples from mildly insulin-resistant C57BL/6J mice, and only 0.65% of the time in samples from highly insulin-sensitive NOD mice (**Table 3**).

**Table 2:**
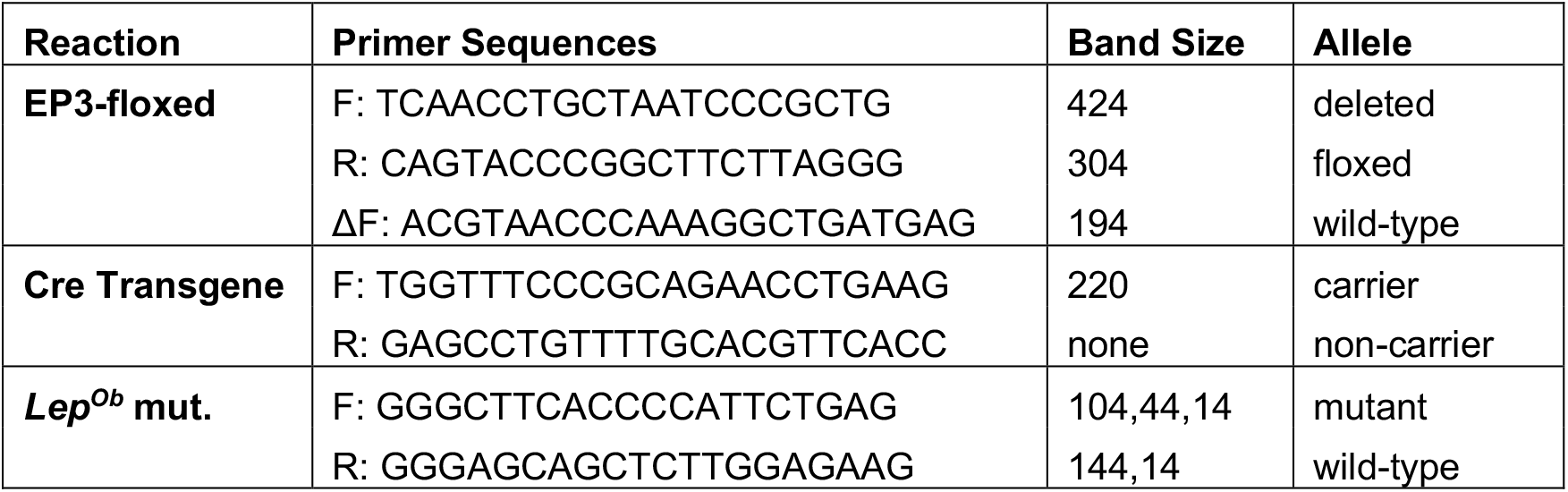
Primer Sequences and expected PCR product sizes for genotyping RIP-Cre-EP3-null lean and *Leptin*^*Ob*^ mice.

**Table 3:**
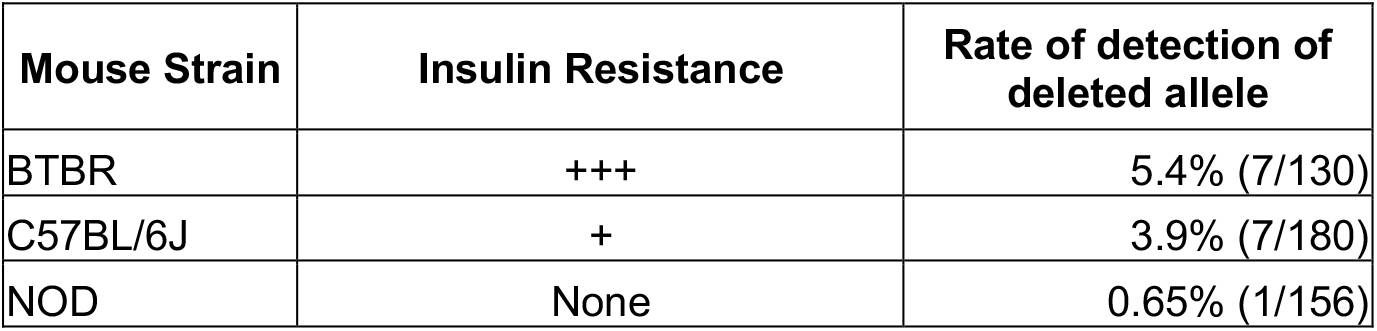
Underlying mouse strain insulin resistance is associated with higher rates of germline transmission of deleted alleles of a different floxed gene in our RIP-Cre^*Herr*^ colonies. The number of reactions used to generate each percentage is indicated in parentheses.

| Mouse Strain | Insulin Resistance | Rate of detection of deleted allele |
| --- | --- | --- |
| BTBR | +++ | 5.4% (7/130) |
| C57BL/6J | + | 3.9% (7/180) |
| NOD | None | 0.65% (1/156) |

**Figure 1.**
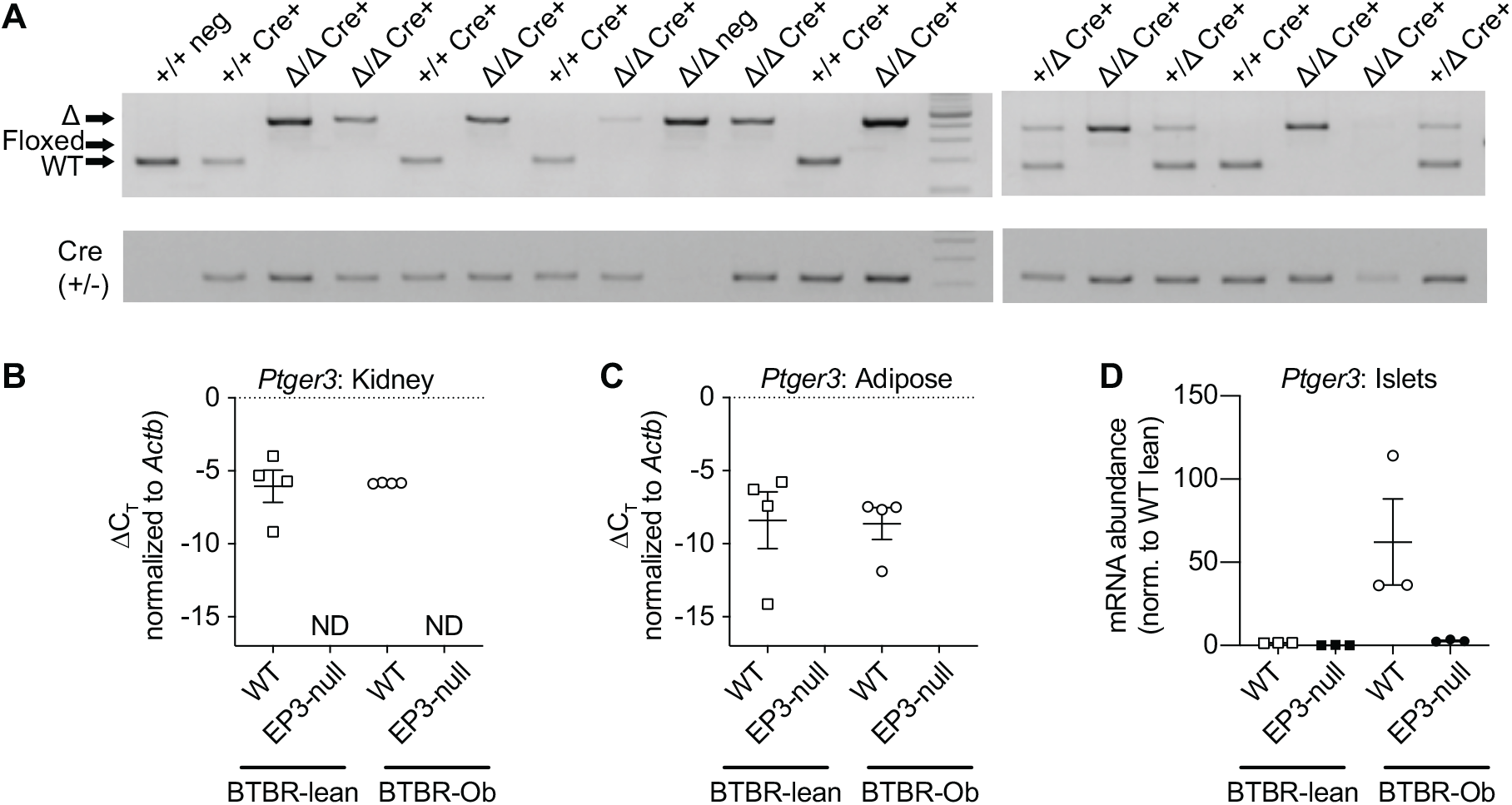
Confirmation of germline transmission of the EP3-floxed deleted allele resulted in a full-body EP3-null BTBR mouse. A: Genotyping results from ear punches showing only the wild-type or deleted allele, and not the floxed allele, in all genotyping reactions, regardless of Cre expression. B and C: qPCR cycle times (C_T_) as normalized to beta-actin (*Actb*) using primers common to all three mouse EP3 splice variants in kidney (B) and adipose tissue (C). ND = no amplification detected. D: qPCR results from islet cDNA samples confirm loss of EP3 expression is nearly or fully complete, with an ablation of EP3 expression in EP3-null BTBR-Ob islets. mRNA abundance was normalized to that of WT via 2^ΔΔCt^ from beta-actin.

### Full-body EP3 loss promotes glucose intolerance and exacerbates insulin resistance in BTBR-lean mice

Body weights and random-fed blood glucose levels were recorded weekly in WT and EP3-null BTBR-lean and -Ob mice from 5-9 weeks of age, with OGTTs performed at 6 weeks of age and ITTs performed at 8 weeks of age. Mice were sacrificed for islet or pancreas collection between 9 and 10 weeks of age.

As expected, BTBR-Ob mice were heavier than BTBR-lean mice at all ages, with no differences in lean or Ob mice by EP3 genotype (**Figure 2A**). Random-fed blood glucose levels were similarly elevated in BTBR-Ob mice compared to BTBR-lean mice at all ages, again, with no difference by EP3 genotype (**Figure 2B**). At 6-8 weeks of age, the 4-6 h fasting blood glucose levels in BTBR-Ob mice were all in the T2D range (> 300 mg/dl), with no differences by EP3 genotype (**Figure 2C**). Because of the lack of protective effect of EP3 loss on BTBR-Ob phenotype, we focused our efforts on characterizing the BTBR-lean mice.

**Figure 2.**
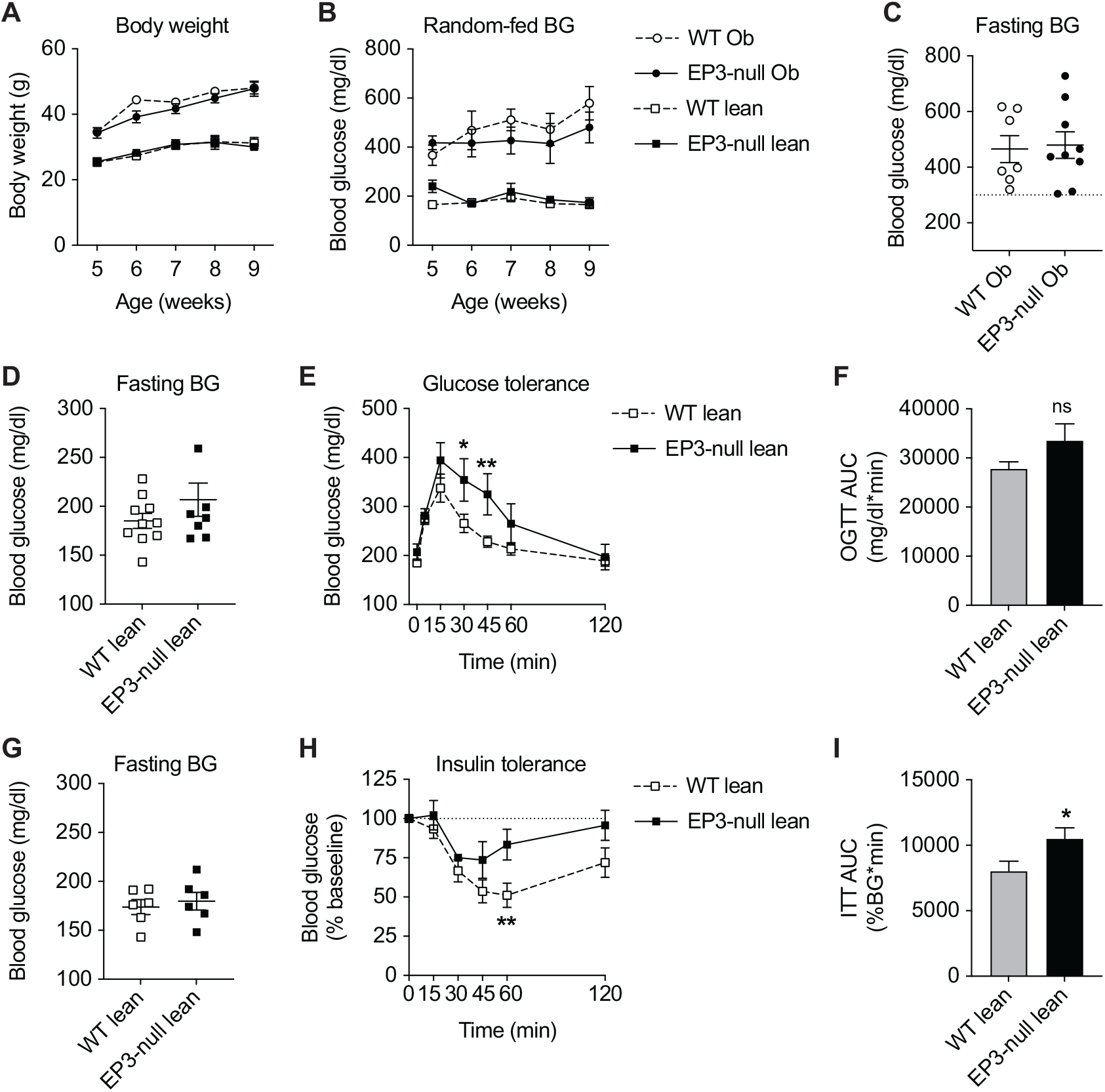
Full-body EP3 loss promotes glucose intolerance and insulin resistance in young, lean BTBR mice. A-B: Body weights (A) and random-fed blood glucose levels (B) of 5-9-week old WT and EP3-null BTBR-lean and BTBR-Ob mice. N=7-11 mice per group. 4-6 h fasting blood glucose levels of 6-week-old WT and EP3-null BTBR-Ob mice. N=7-8 mice per group. D-F: 4-6 h fasting blood glucose levels (D), blood glucose levels after oral glucose challenge (E) and OGTT AUCs (E) of 6-week-old WT and EP-null BTBR-lean mice. N=7-9 mice per group. G-I: 4-6 h fasting blood glucose levels (G), reduction in blood glucose levels after IP insulin injection (H) and ITT AUCs (I) of 8-week-old WT and EP-null BTBR-lean mice. N=6 mice per group. In A, B, E, and H, data were compared between genotypes by two-way ANOVA with Holm-Sidak test post-hoc to correct for multiple comparisons. in C, D, F, G, and I, data were compared by two-tail T-test. ^*^, p < 0.05 and ^**^, p < 0.01 vs. control. ns = not significant. If no p-value is indicated, the difference in means was not statistically significant.

At 6 weeks of age, there was no statistically significant difference in the mean 4-6 h fasting blood glucose level of EP3-null and WT BTBR-lean mice by genotype (**Figure 2D**), although EP3-null BTBR-lean mice were significantly more glucose intolerant than WT controls (**Figure 2E,F)**. At 8 weeks of age, there was still no difference in mean 4-6 h fasting blood glucose levels between WT and EP3-null BTBR lean mice (**Figure 2G**), but, when given an insulin challenge, EP3-null BTBR-lean mice were significantly less insulin sensitive than WT controls (**Figure 2H,I**).

### Full-body EP3 loss does not affect markers of beta-cell replication in BTBR-lean islets and pancreas sections

At 9-10 weeks of age, pancreases were collected from WT and EP3-null BTBR-lean mice and fixed for paraffin-embedding and sectioning. Insulin IHC was performed on pancreas sections counterstained with hematoxylin (see representative images in **Figures 3A and 3B**). There was no difference in beta-cell fractional area between WT and EP3-null BTBR-lean pancreas sections (**Figure 3C**). Co-immunofluorescence experiments for insulin and nuclear Ki67 revealed no change in actively-replicating beta-cells by genotype (**Figure 3D**). These findings are contradictory to those of a published study showing islets from high fat-diet fed EP3-null C57BL/6J mice have significantly elevated beta-cell replication (24). Therefore, qPCR for Cyclin D1 (gene symbol: *Ccnd1*), whose expression is required for cell cycle progression, was performed on islet samples from 10-week-old WT and EP3-null BTBR mice as compared to C57BL/6J. BTBR islet *Ccnd1* expression was unchanged by genotype, but was nearly 12-fold lower than that of C57BL/6J islets, consistent with an intrinsic block in beta-cell replication as a BTBR strain characteristic (5-7) (**Figure 3E**).

**Figure 3:**
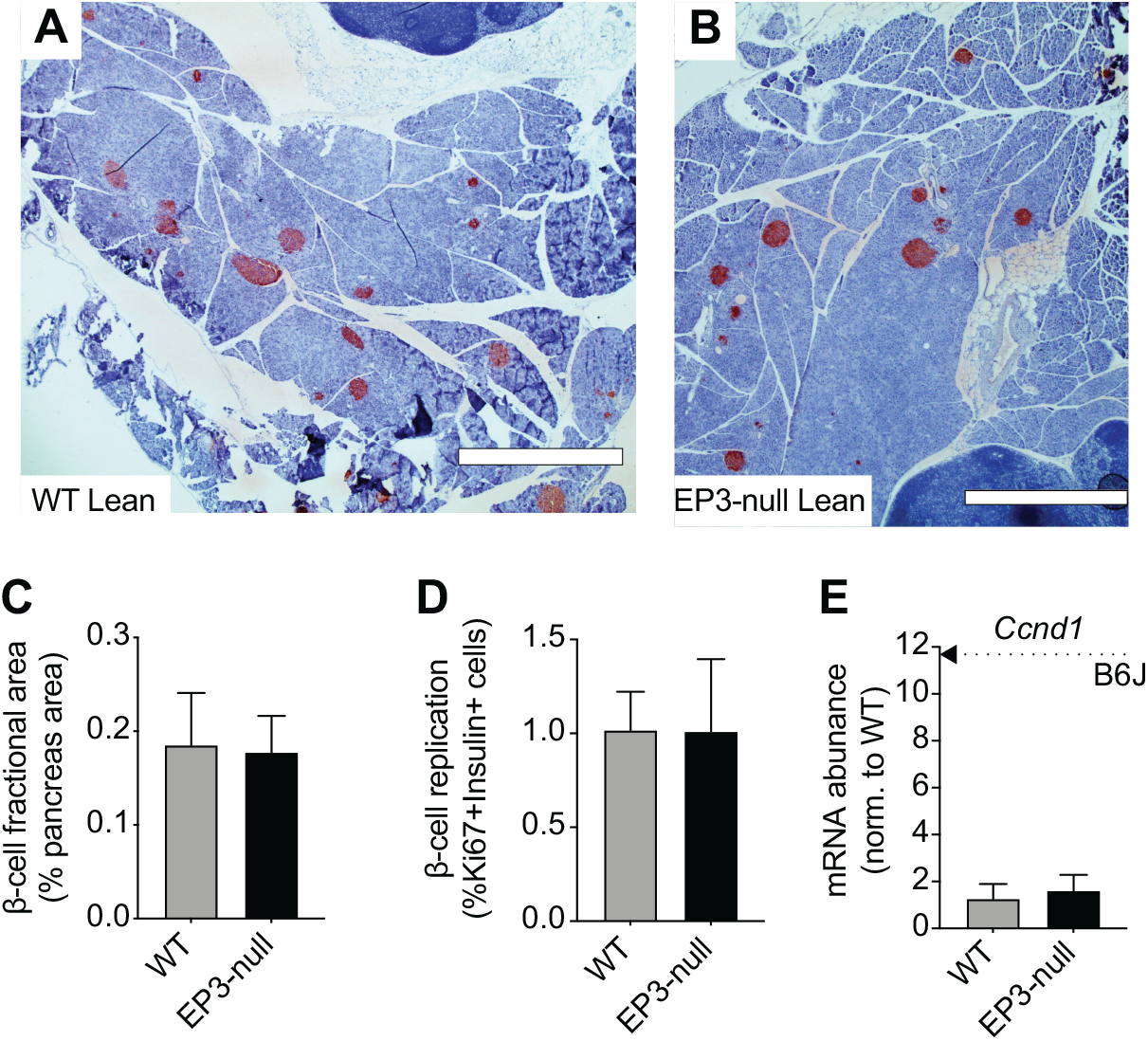
The EP3-null mutation has no effect on BTBR-lean β-cell fractional area or markers of β-cell proliferation. A-B: Representative insulin immunohistochemistry images from WT BTBR-lean mice (A) and EP3-null BTBR-lean mice (B). Scale bars = 1000 µm. C: Quantification of insulin-positive area from the immunostaining shown in (A). N=3 mice per group, with at least 2 pancreas sections analyzed per mouse. D: Quantification of actively replicating beta-cells as determined by co-immunofluorescence of slide sections for Ki67 and insulin. As in (C), N=3 mice per group, with at least 2 pancreas sections analyzed per mouse. E: Cyclin D1 (*Ccnd1*) mRNA abundance in WT and EP3-null BTBR-lean islets as compared to the mean of C57BL/6J (B6J) islets (indicated by dashed arrow). mRNA abundance was normalized to that of WT via 2^ΔΔCt^ from beta-actin. N=4-8 mice per group. In all panels, data were compared by two-tail T-test, with none of the differences being statistically significant.

### EP3 loss promotes unregulated glucose-stimulated insulin secretion in BTBR-lean islets, consistent with elevated expression of constitutively-active EP3 variants as a strain characteristic

*Ex vivo* glucose-stimulated insulin secretion (GSIS) assays were performed on islets isolated from 9-10-week old WT and EP3-null BTBR-lean mice in response to 2.8 mM glucose or 16.7 mM glucose, the latter with or without the addition of 10 nM of exendin-4 (Ex-4), a glucagon-like peptide 1 receptor (GLP1R) agonist. The most striking findings from these assays was the complete loss of regulated insulin secretion in islets from EP3-null BTBR-lean mice, with high and nearly equivalent levels of insulin secreted in basal or stimulatory glucose, and no additional effect of Ex4 (**Figure 4A**). This difference in secretion was not due to increased insulin content, as EP3-null BTBR-lean islets had significantly lower insulin content than WT controls (**Figure 4B**).

**Figure 4:**
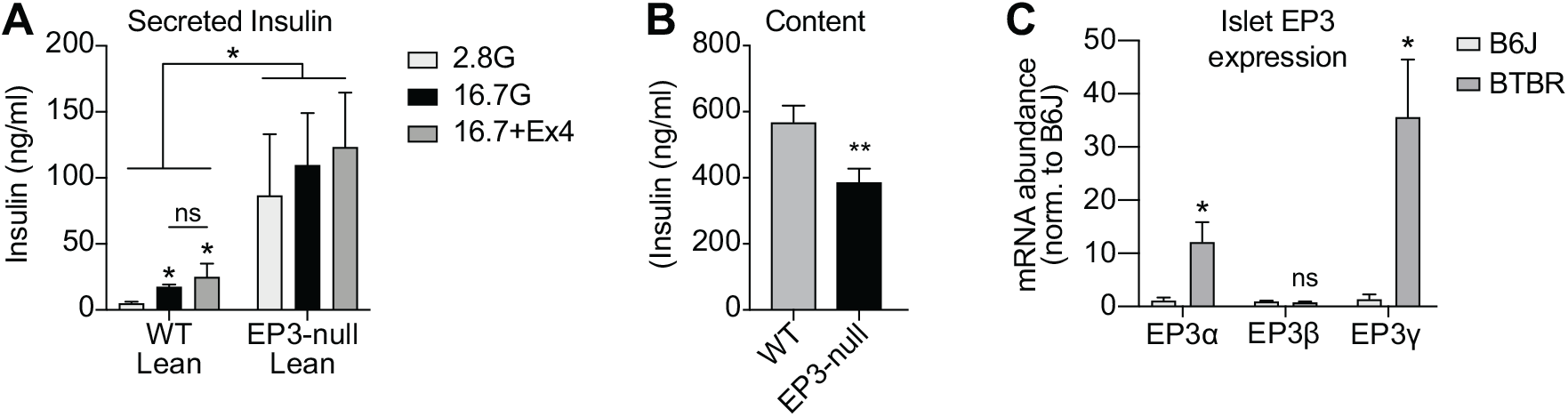
Islets from EP3-null mice hypersecrete insulin *ex vivo*, consistent with elevated islet expression of constitutively-active EP3 splice variants. A. Insulin secreted over 45 min from WT and EP3-null BTBR-lean islets in response to 2.8 mM glucose, 16.7 mM glucose, or 16.7 mM glucose + 10 nM exendin-4 (Ex4). N=6-11 mice/group. B: Islet insulin content from the experiments shown in (A). C: EP3 splice variant abundance in islet cDNA samples from 10-week-old C57BL/6J (B6J) or BTBR-lean males using primers specific for each of the mouse EP3 splice variants. mRNA abundance was normalized to that of B6J islets via 2^ΔΔCt^ from beta-actin. N=4-8 mice per group. In A, data were compared between genotypes by two-way ANOVA and within genotype by one-way ANOVA with Holm-Sidak test post-hoc to correct for multiple comparisons. In B and D, data were compared between genotypes by two-tail T-test. ^*^, p < 0.05 and ^**^, p < 0.01 vs. control. ns = not significant. If no p-value is shown, the results were not statistically significant.

A previous report had shown no impact of full-body EP3 loss on C57BL/6J islet GSIS (24). In order to investigate a root cause of these strain-specific effects, qPCR was performed on islet cDNA samples from 10-week-old BTBR or C57BL/6J mice with primers specific for each of the three mouse EP3 splice variants (EP3α, EP3β, EP3γ). EP3α and EP3γ mRNA abundance was ∼12-fold and 36-fold higher, respectively, in islets from BTBR-lean mice as compared to C57BL/6J, with EP3β mRNA abundance being unchanged (**Figure 4C**). Notably, EP3α and EP3γ have partial to near-full constitutive activity, respectively, meaning no ligand is necessary in order for these variants to signal to downstream effectors (12, 14, 18, 38, 39).

### Differences in islet EP3 expression between Balb/c males and females explain the sexually-dimorphic phenotype of Gα_z_-null Balb/c mice

Gα_z_ is an inhibitory G_i/o_ subfamily member that exclusively mediates the inhibitory effect of EP3 on beta-cell cAMP production and GSIS (10-12, 14). Previous work from our laboratory revealed 9-12-week-old female Balb/c mice deficient in Gα_z_had reduced fasting blood glucose levels and accelerated glucose clearance after glucose challenge (33). With the extremely restricted tissue distribution of Gα_z_and significantly increased plasma insulin levels during GTTs, this phenotype was best explained by a beta-cell-centric mechanism (10, 33, 40, 41). Like females, 9-12-week old male WT and Gα_z_-null Balb/c mice were of equivalent weight as WT controls (**Figure 5A**). Unlike females, though, the 4-6 h fasting blood glucose levels of male Gα_z_-null Balb/c mice were unchanged as compared to controls (**Figure 5B,C**). Also unlike females, there was no difference in male Gα_z_-null Balb/c glucose tolerance (**Figure 5D,E**). Consistent with their *in vivo* phenotype, but in contrast to results from female mice, there was no increase in plasma insulin of Gα_z_-null males after glucose challenge (**Figure 5F**). Upon qPCR analysis of male and female Balb/c islet cDNA samples, all three EP3 splice variants were significantly enriched in islets from female Balb/c mice as compared to male, with EP3γ being the most highly so (∼10-fold increased) (**Figure 5F**). Finally, there was no impact of Gα_z_loss on male or female Balb/c mouse insulin sensitivity, although male Balb/c mice were almost completely insulin resistant as compared to highly insulin-sensitive females (**Figure 5G**).

**Figure 5.**
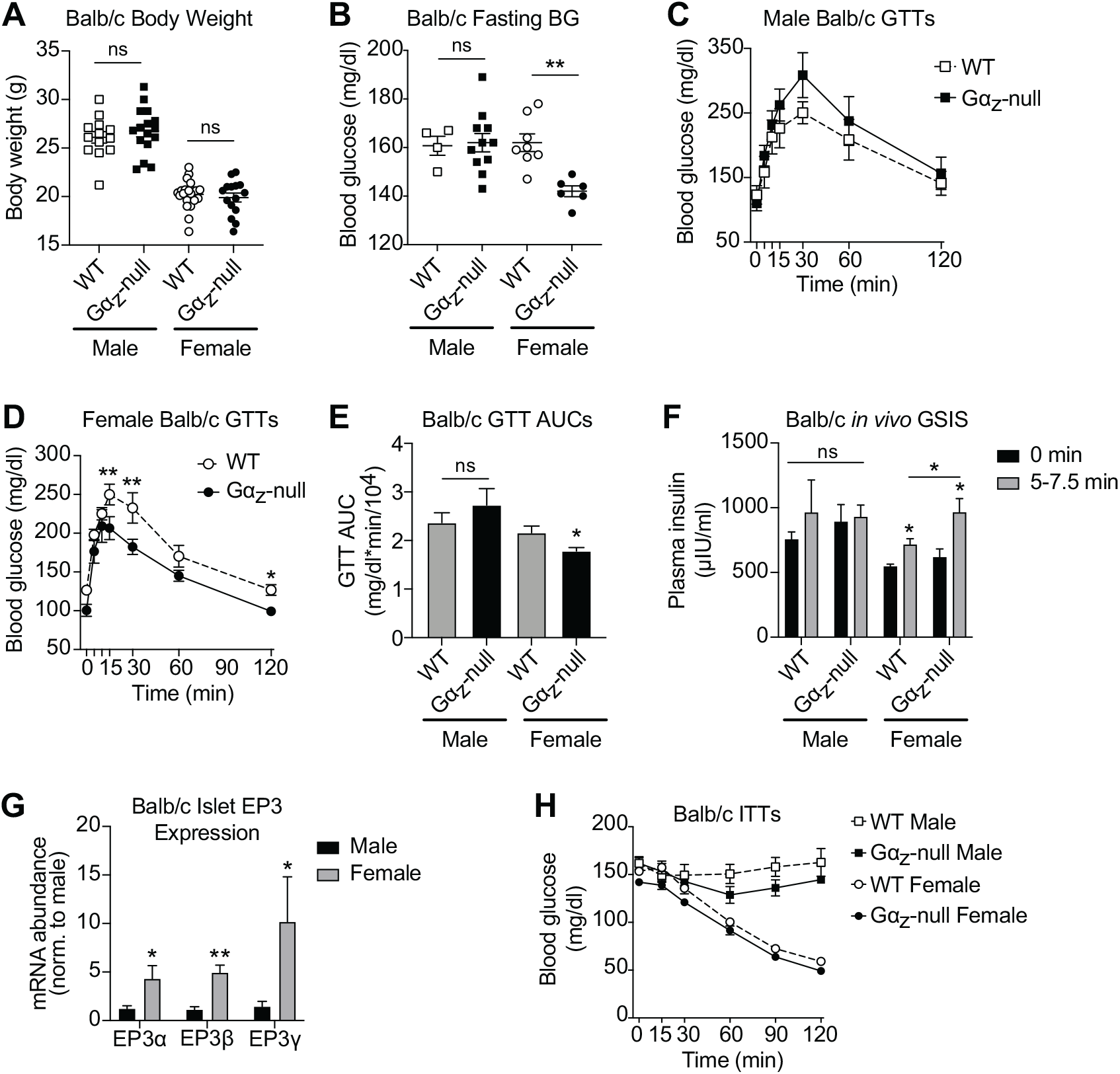
Differential expression of EP3 splice variants explains the sexually-dimorphic metabolic phenotype of male and female Gα_z_-null Balb/c mice. A-B: Body weights (A) and 4-6 h fasting blood glucose levels (B) of 9-12-week old male and female WT and Gα_z_-null Balb/c mice. C-D: Blood glucose levels after IP glucose challenge of WT and Gα_z_-null male Balb/c mice (C) and female Balb/c mice (D). N=5-12 mice per group. E: Area-under-the curve (AUC) analyses of the data shown in (C) and (D). F: Plasma insulin levels before and 5-7.5 min after glucose challenge during the GTTs shown in (C) and (D). G: qPCR of islet cDNA samples from 10-week-old male and female Balb/c mice for each of the mouse EP3 splice variants, with mRNA abundance normalized to that of male islets via 2^ΔΔCt^ from beta-actin. N=4-6 mice per group. H: Blood glucose levels of 11-12-week old male and female WT and Gα_z_-null Balb/c mice after IP insulin administration. N=3-12 mice per group. In panels A, D, E, F, and H, data from female mice are adapted from (33) under the terms of the Creative Commons CC-BY license, which permits unrestricted use, distribution, and reproduction in any medium, provided the original work is properly cited. In A, B, G, and E, data were analyzed within sex by two-tail T-test. In C, D, F, and H, data were analyzed within sex by two-way ANOVA with Holm-Sidak test post-hoc to correct for multiple comparisons. ^*^, p < 0.05 and ^**^, p < 0.01 vs control. ns = not significant. If no p-value is shown, the results were not statistically significant.

## Discussion

We and others have previously proposed elevation in islet PGE_2_ production and EP3 expression as a dysfunctional response to the T2D condition, actively contributing to the loss of functional beta-cell mass in T2D (8, 9, 13, 17, 18, 20, 23, 42). On the other hand, previous studies found global EP3 loss accelerated the development of insulin resistance and/or glucose intolerance in diet-induced obesity (DIO) models (24-26) and that systemic administration of an EP3-specific antagonist had no effect on the T2D phenotype of DIO-induced C57BL/6J mice (27). In the T2D BTBR-Ob model, treatment of islets *ex vivo* with an EP3 antagonist or reducing PGE_2_ production via a dietary intervention improves their insulin secretion response to glucose and GLP1R agonists, highlighting the importance of EP3 signaling in their beta-cell dysfunction (8, 9). In this work, we aimed to test the hypothesis beta-cell-specific loss of EP3 would protect BTBR-Ob mice from developing T2D by allowing a beta-cell compensation response to occur. Ultimately, our mouse model was confirmed as a full-body EP3 knockout, and, like the DIO models, full-body EP3 loss did not confer any protection to BTBR-Ob mice from T2D. Unexpected based on the preponderance of literature were our findings that lean, EP3-null BTBR mice developed early and prominent insulin resistance, glucose intolerance, and islet insulin hypersecretion – phenotypes previously unobserved in lean EP3-null C57BL/6J mice (24, 43). Finally, for the first time, the direct correlation of lean mouse islet expression of constitutively-active EP3 variants in a strain- and/or sex-specific manner with the effect of EP3 or EP3-coupled Gα_z_ loss on metabolic phenotype implicates the EP3 signaling pathway as a necessary, functional regulator of insulin secretion and not simply a dysfunctional consequence of T2D.

In the current work, we employed a three-primer genotyping strategy to identify the *Ptger3* deleted allele in ear tissue, suggesting germline recombination in the founder mouse had created a full-body knockout – a finding confirmed by qPCR of adipose, kidney, and islet tissue. In searching the literature, we found a previous report using the RIP-Cre^*Herr*^ line in a mixed B6/CBAJ background, in which a three-primer genotyping reaction revealed the deleted allele for a protein-of-interest was detected in < 6% of tail tissue from F2 progeny mice (44). As in the current work, Spinelli and colleagues found that when the deleted allele was detected in the tissue harvested for genotyping, the mouse was a full-body knockout (44). Additional studies by Spinelli and colleagues revealed absence of the deleted allele in the genotyping reaction was essentially a surrogate of cell-specific deletion, providing a simple screening mechanism to ensure the utility of the RIP-Cre^*Herr*^ driver line (44). While Spinelli and colleagues did not speculate on the mechanisms behind this recombination, the EP3-Flox-RIP-Cre line was re-derived by *in vitro* fertilization of wild-type BTBR female oocytes with transgenic male sperm, meaning germline recombination must have happened in the spermatocytes. Our novel finding the prevalence of germline deleted allele transmission with the RIP-Cre^*Herr*^ driver line is directly related to the underlying insulin resistance of the background strain suggests a natural phenomenon based on physiologically-relevant activation of the insulin gene promoter, and not simply ectopic gene expression. Interestingly, there is a significant body of literature on the role of cell-autonomous insulin gene expression, secretion, and signaling in spermatocyte maturation and function. Insulin protein is expressed and in isolated mammalian spermatocytes, and insulin is secreted *ex vivo* in a pulsatile, glucose-dependent manner (45, 46). Insulin signaling induces pig sperm capacitation and acrosome reaction, and blocking endogenous sperm insulin exocytosis using the calcium channel blocker, nifedipine, ablates these effects (46). Both insulin and leptin regulate human sperm glucose metabolism and glycogen synthesis in an autocrine manner, which Aquilla and colleagues speculate allows sperm to survive and function completely autonomously in transit to the oocyte(45, 47). Yet, BTBR males have high motile sperm counts, and BTBR mice are considered exceptional breeders (32). In consideration of the underlying insulin resistance of the BTBR strain, it is plausible their spermatocytes have to produce and secrete more insulin to compensate for this insulin resistance, just as their beta-cells do, making a recombination event more likely than in an insulin-sensitive strain.

Four previous studies have metabolically phenotyped the effect of full-body EP3 loss or EP3 blockade in the context of C57BL/6J diabetogenic mouse models (24-27). Sanchez-Alavez and colleagues maintained wild-type and EP3-null C57BL/6J mice on breeder chow (approximately 20 kcal% fat, as compared to 10 kcal% fat in normal rodent chow) for 40 weeks, revealing EP3-null mice gradually became heavier than wild-type controls, developing hyperinsulinemia, hyperleptinemia, and worsening glucose tolerance between 3 and 6 months of age (25). Similar results were found in breeder chow-fed EP3-null C57BL/6J mice by Xu and colleagues, with the additional conclusion that constitutively-active EP3α and EP3γ are the primary limiters of adipose tissue lipolysis (26). Ceddia and colleagues used a 16-week 45 kcal% fat high-fat diet (HFD) regimen, initiated at 4 weeks of age in WT and EP3-null C57BL/6J mice, revealing a diet-dependent decrement in insulin sensitivity correlated with unregulated lipolysis, dyslipidemia, and systemic inflammation secondary to EP3 loss in adipose tissue(24). At 20 weeks of age, HFD-fed EP3-null mice were significantly less glucose tolerant than their wild-type counterparts(24). A second study by Ceddia and colleagues tested the effect of 1 week of systemic treatment with the EP3 antagonist, DG-041, on the metabolic phenotype of 16-week HFD-fed C57BL/6J mice, with the addition of a 60 kcal% fat-fed (very HFD: VHFD) group (27). While few-to-no gross changes in full-body metabolic phenotype were observed, DG-041-adminstered VHFD-fed mice had higher liver triglyceride content and more pronounced hepatic steatosis, suggestive of exacerbated liver insulin resistance (27). A secondary, strong conclusion from results of these studies, though, as well as a fifth studying the effect of full-body EP3 loss on C57BL/6J kidney water resorption (43), is systemic EP3 loss is insufficient to induce metabolic dysfunction in lean C57BL/6J mice fed standard, low-fat, chow or semi-purified control diets. Because of this, we expected no phenotype of EP3 loss in young, lean BTBR males, intending to simply use them as controls for the BTBR-Ob study. If the underlying insulin resistance phenotype of the BTBR strain were exacerbated solely by loss of adipose EP3, the rapid and significant effect of full-body EP3 deletion on BTBR-lean mouse glucose tolerance would not be unexpected. Based our results with female Gα_z_-null Balb/c mice, though, which remain as highly insulin sensitive as WT controls, we suspect beta-cell EP3 loss and insulin hypersecretion, in the context of underlying insulin resistance, is required for the full metabolic phenotype of EP3-null BTBR-lean mice.

We suspect, but cannot confirm in the short time course study presented in the current work, EP3-null BTBR-lean mice would ultimately have progressed to T2D. BTBR islets secrete less insulin in response to glucose or GLP1R agonists than those from the commonly-used C57BL/6J strain (8). This phenotype has been linked to a sNP in the Tomosyn-2 gene that interferes with insulin granule exocytosis (3, 4). While 5-9-week-old EP3-null BTBR-lean mice were clearly more insulin resistant and glucose intolerant than WT controls, they did not have elevated fasting blood glucose levels: a key component to T2D. Prominent and abnormally-regulated insulin hypersecretion, with no impact on beta-cell replication or fractional area, indicates EP3-null beta-cells from BTBR-lean mice are under a great deal of stress: a condition that should eventually cause beta-cell failure. This concept has support in epidemiological studies in human populations with an underlying genetic defect predisposing to skeletal muscle insulin resistance, in which the progression to T2D is dependent on the beta-cell (48-52). Future studies will be necessary to tease apart the timeline of metabolic dysfunction in the BTBR line after EP3 loss.

Beta-cell EP3 has been shown in numerous studies to couple specifically to the unique inhibitory G protein, G_z_, to reduce cAMP production and GSIS (10-12, 34). C57BL/6N or 6J mice lacking Gα_z_ throughout the body or specifically in the beta-cells, respectively, are completely protected from HFD-induced T2D, with no effect on body weight or systemic insulin resistance (11, 12).

In this study, we found expression of EP3α and EP3γ, splice variants with partial-to-near-full constitutive activity, were significantly up-regulated in islets BTBR-lean mice as compared to C57BL/6J. Notably, EP3γ has an absolute requirement for beta-cell Gα_z_ in order to inhibit cAMP production and GSIS, and it does so in the complete absence of ligand (12). In the BTBR strain, the purpose of up-regulation of constitutive EP3 activity and whether it is functional or dysfunctional remains undetermined. Strong support for a functional up-regulation comes from the fact EP3γ expression is also significantly up-regulated in islets from highly insulin-sensitive female Balb/c mice. When deficient in Gα_z_, female Balb/c mice display prominent fasting hypoglycemia, accelerated glucose clearance, and insulin hypersecretion *in vivo* (33). An intriguing hypothesis is high constitutive beta-cell EP3 activity is required in conditions where a strong check on insulin secretion is required. In female Balb/c mice, this could be a mechanism to prevent hypoglycemia in the context of high insulin sensitivity. In BTBR mice, this could be a mechanism to prevent beta-cell failure in the context of beta-cell stress secondary to underlying defects in skeletal muscle insulin resistance and insulin exocytosis.

In sum, using two different lean mouse strains, the current work is the first description of a critical role for endogenous islet EP3 signaling in regulating systemic glucose homeostasis and metabolic function in the absence of diabetes. Considered in the context of the existing body of literature, including recent work from our own laboratory associating up-regulated islet EP3 signaling with functional beta-cell compensation in non-diabetic obese humans (15), these results have significant implications for the study of EP3 as a therapeutic target for the prevention or treatment of beta-cell dysfunction and T2D.

## Acknowledgments

We wish to thank the many present and former members of the Kimple Laboratory who contributed technical assistance or scientific discussion during the course of these experiments. A particular note of thanks goes to Audrianna Wu, a high-school student who completed a summer research project on the EP3-null mouse phenotype. We also wish to thank Patrick J. Casey for his contributions to and scientific discussion of the Balb/c phenotype. This work was supported in part by Merit Review Award I01 BX003700 from the United States (U.S.) Department of Veterans Affairs Biomedical Laboratory Research and Development Service (BLR&D) (to MEK). Further support was provided by NIH Grants K01 DK080845 (to MEK), R01 DK102598 (to MEK), F31 DK109698 (to RJF), and R01 DK076488 (to PJC); ADA Grant 1-16-IBS-212 (to MEK), and JDRF Grant 17-2011-608 (to MEK).

## Author Contributions

Conceptualization, JAW and MEK; data curation, AR, JAW, and MEK; formal analysis, AR, JAW, and MEK; funding acquisition, RJF and MEK; investigation, JAW, AR, DP, MDS, and RJF; methodology, JAW, AR, and MEK; project administration, MEK; supervision, JAW and MEK; visualization, JAW, AR, and MEK; writing—original draft, JAW, AR, and MEK; writing—review and editing, JAW and MEK. All authors have read and agreed to the published version of the manuscript.

